# Kiosc: an integrated platform for managing bioinformatics data analysis containers

**DOI:** 10.64898/2026.08.20.745983

**Authors:** Federico Marotta, Oliver Stolpe, Benedikt Obermayer, January Weiner, Manuel Holtgrewe, Dieter Beule, Mikko Nieminen

## Abstract

**Background:** In many bioinformatic data analysis projects, it is convenient to visualize plots and results through an interactive web app or dashboard. These interactive reports can then be shared with customers, collaborators, or the general public. Publishing and sharing these apps is not straightforward, becoming especially cumbersome when the number of projects and customers start growing.

**Methods:** Docker containers offer a convenient way to package, distribute, and run interactive web apps, and their use is already widespread in the bioinformatics community. We developed Kiosc to simplify the orchestration of containerized web apps, organize them into projects, and regulate access control. We implemented it as a web server based on the Django framework, with a user- and admin-friendly interface as well as a REST API for programmatic tasks. Users can select Docker containers packaging apps like Plotly Dash, Shiny, or Quarto, and configure them to display the results of their analysis. Kiosc runs the containers with the appropriate network configuration and acts as a proxy to the web services running inside the containers.

**Results:** We have been maintaining a Kiosc instance for more than 5 years, serving 321 containers in 150 projects across multiple institutions. In this article, we introduce the main functionality in Kiosc and describe four use-cases that show how Kiosc can prove helpful to the broader bioinformatics community, such as configuring and running web apps for the interactive visualization of workflow results, and publishing companion apps for scientific articles.

**Conclusion:** Kiosc is a self-hosted platform for publishing web apps, which doesn’t require significant expertise in either Docker or network administration to be deployed. It provides a similar service to Kubernetes, but with a convenient web interface and much lower administration overhead.

## Introduction

Data exploration and results visualization are integral parts of modern bioinformatic workflows (Holmes & Huber, 2019). Both these tasks can be performed through interactive web applications (apps), which offer several advantages over both command-line tools and static PowerPoint presentations. For example, the viewers of a web app can navigate directly to plots and tables resulting from an analysis, change the input parameters through widgets, and in some cases perform the same analysis on their own data. Furthermore, these apps do not have to be installed and are therefore highly portable across operating systems and devices, requiring only a web browser to be accessed. Interactive web apps increase the reproducibility of an analysis and make new methods available even to users with no programming experience. Sharing these apps with customers and collaborators establishes a bridge between the analyst or method developer and the biomedical scientist with little experience in computational and statistical frameworks. In recent years, more and more bioinformatics tools are made available as web apps, rather than just as command-line tools (CZI Cell Science Program, 2025; Gao et al., 2013; McGowan et al., 2020; Obermayer et al., 2020; Speir et al., 2021; van Kempen et al., 2024). Libraries like Shiny (Chang et al., 2024), Plotly Dash (https://plotly.com), Streamlit (https://streamlit.io), and Jupyter (Granger & Perez, 2021) make it easy to develop such web apps for bioinformatics.

Bioinformatics groups and institutions often want to use, build, and publish a large number of web apps. Modern development ecosystems have the drawback of requiring a large number of additional operating system libraries and software dependencies to build and run most apps, making it challenging to run multiple apps on a single machine. Virtualization and containerization technologies emerged as an attempt to simplify building and deploying software stacks and were quickly adopted by the scientific community as a way to boost the reproducibility of research (Boettiger, 2015; Kulkarni et al., 2018; Nüst et al., 2020). One technology in particular, Docker, became the *de facto* standard for containerized apps (Madhavapeddy et al., 2026, https://docker.com), and is now supported by all major workflow managers (Di Tommaso et al., 2017; Galaxy Community, 2024; Mölder et al., 2021). While Docker on its own simplifies deployment of web apps, setting up an environment with multiple containers can require a high level of technical expertise. Containerized web apps introduce the additional challenge of requiring network configuration, persistent data storage, and access control. Thus, hosting bioinformatics web apps is still a significant challenge.

The companies behind Shiny, Plotly Dash, and other popular web app frameworks typically offer paid or freemium services for hosting apps, but these products are limited to apps implemented in the respective programming language (https://plotly.com/cloud; https://posit.co/products/enterprise/connect-cloud;https://posit.co/products/open-source/shiny-server; https://streamlit.io/cloud). Therefore, some popular apps such as cBioPortal (Gao et al., 2013) and CELLxGENE (CZI Cell Science Program, 2025), which are not implemented in any of these frameworks, would be left out. ShinyProxy (https://www.shinyproxy.io) is another flexible option for hosting data science apps, supporting access control and scalability of resources. However, adding new apps entails modifying the configuration file and restarting ShinyProxy itself; although it is possible to avoid downtime by using ShinyProxyOperator, the task of adding or updating apps is centralized in the configuration file and is typically relegated to the administrator. The current industry-grade standard for orchestrating general-purpose containers is Kubernetes (https://kubernetes.io), which offers advanced features but has a steep learning curve and requires significant computational and human resources for its maintenance (Shamim et al., 2022). Other existing solutions, either commercial or made for “home labs”, such as Portainer (https://www.portainer.io) and Dockge (https://dockge.kuma.pet), lack important features in a multi-user setting such as role-based access control, at least in the free community edition.

We developed Kiosc (Kiosc is Our Support for Containers) as an intermediate solution that allows hosting and managing Docker containers in simple settings, while still satisfying the needs of bioinformatics groups who want to host and granularly share multiple web apps. The focus of Kiosc is specifically on containerized web apps for interactive data exploration, results visualization, or teaching purposes, as opposed to heavy-duty analysis of big data. Other tools like Galaxy, Snakemake, or Nextflow are better suited to run analysis workflows in a high-performance computing setting, since they allow to scale the resources assigned to each container and can connect the output of a container to the input of the next, thus building a proper workflow. Kiosc, on the other hand, offers convenience tools for web apps, such as access control, persistent data storage, proxying of incoming requests to the various web services, and the ability for users to dynamically and autonomously add new apps, all through a unified web interface.

## Methods

### Implementation

Kiosc itself is a web app implemented on the SODAR Core framework, a Django-based system for scientific data management (Django, 2025; Nieminen et al., 2020). SODAR Core provides authentication, project management, project-specific access control, asynchronous background jobs support, and a storage system for small files. On top of this, Kiosc adds container-specific functionality, such as the ability to pull Docker images from registries, start and stop the containers, and access their logs. All the operations can be carried out from the web user interface, and a separate REST API provides endpoints for programmatically retrieving container data and automating some tasks. Furthermore, Kiosc acts as a reverse proxy, abstracting away the need for port management and providing a unified web interface to all web apps for the user (Figure 1A).

**Figure 1.**
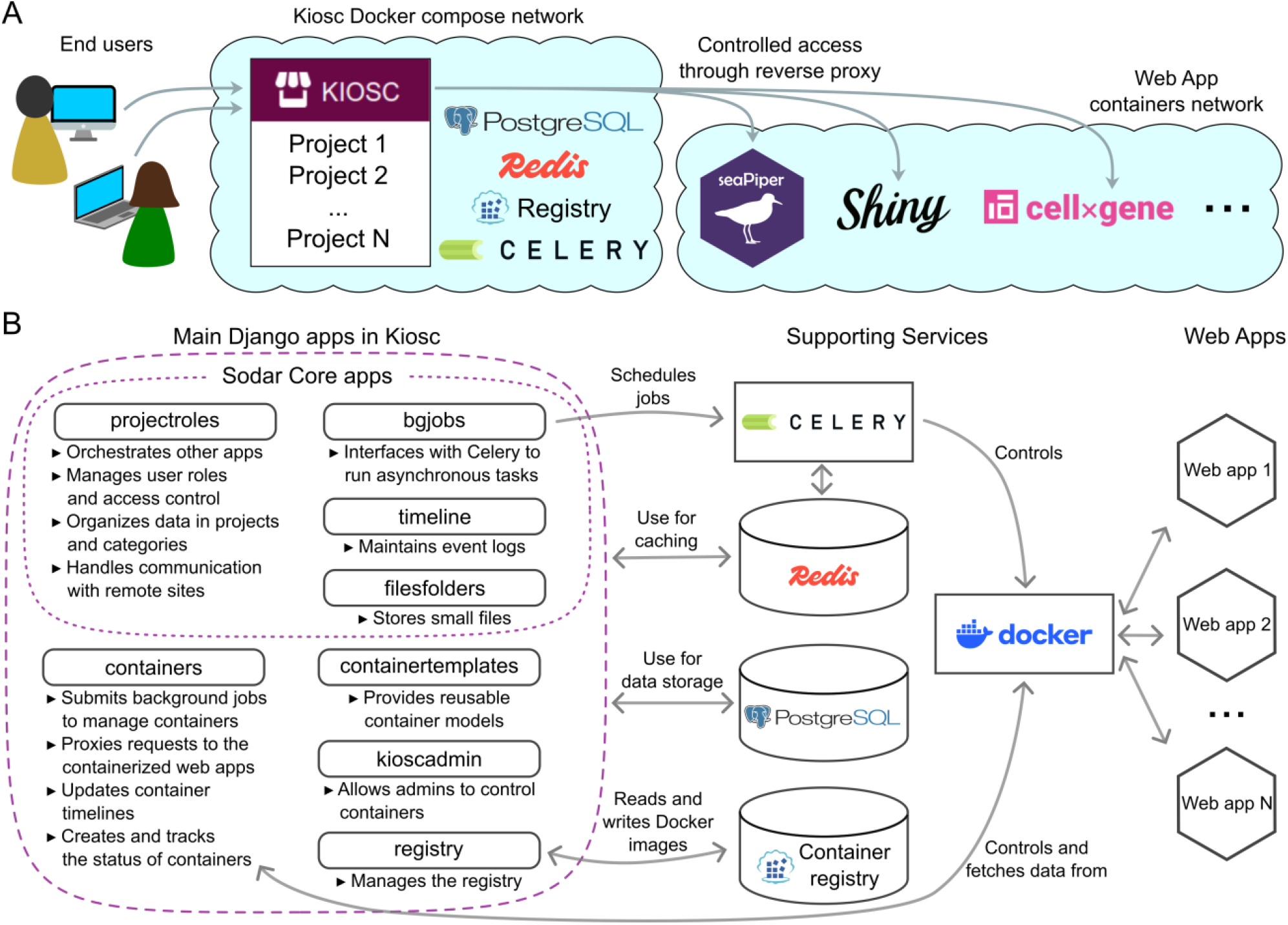
Kiosc architecture. (A) End users connect to the web interface, where they have access to the list of projects. Kiosc is built as a Django server supported by PostgreSQL, Redis, Celery, and a Container Distribution Registry. In our production environment, these services run in a Docker compose network which is separate from the Docker network where the app containers are launched. Kiosc forwards user requests to the app containers, acting as a reverse proxy. (B) The main Django apps in Kiosc and their relationships to the other components of the software stack.

More specifically, the bulk of Kiosc consists of eight main Django apps which interface with the Docker daemon and additional services (Figure 1B). Four Django apps, `projectroles`, `timeline`, `filesfolders` and `bgjobs`, are part of the SODAR Core framework and provide common functionality which can be shared across websites. The `projectroles` app, which manages user roles and projects, also acts as an orchestrator for the other Django apps. The `filesfolders` and `bgjobs` apps provide storage for small files and asynchronous tasks, respectively, while the timeline app takes care of logging events to a database. The `containers` Django app bridges Kiosc to the underlying Docker containers by representing each container as an entry in the database, and by using the `docker-py` Python library to control the containers. Time-consuming operations on containers are performed asynchronously through Celery workers to avoid blocking the website, a functionality offered by the bgjobs app. The `containers` Django app also implements the reverse proxy through which the web apps inside the containers are accessed. A separate Django app, `kioscadmin`, can be used by site administrators to check all running containers and perform Docker-related tasks from the web interface. This app also defines periodic background jobs to stop containers after inactivity and to synchronize the state of a container to its database representation, so that Kiosc can provide up-to-date information to the user even when the container crashes. The `containertemplates` app maintains models which can be reused across projects, so that users don’t have to fill all the fields of the form when creating a container. Finally, the `registry` Django app interfaces with the container registry integrated with Kiosc. The registry is subject to the same authentication and authorization rules as the main Kiosc site, and once an image is pushed to it, a corresponding container object is made available to the user automatically.

Besides Django and Python dependencies, Kiosc relies on additional services which must be available at runtime (Figure 1B). The PostgreSQL database (PostgreSQL, 2026) stores data about users, projects, containers, and event logs. The Celery daemon (https://docs.celeryq.dev) schedules periodic clean-up tasks and runs user-submitted background jobs to control the Docker daemon. Redis (https://redis.io) is used by Django for caching and by Celery for brokering and storing data about the background jobs. Furthermore, it is used by a Django Channels websocket (https://channels.readthedocs.io) to receive and stream container status updates and logs in real time. A self-hosted CNCF Container Distribution Registry (https://distribution.github.io/distribution) stores Docker images which can only be accessed through Kiosc, hence it is useful as a private space for containers which contain sensitive information and shouldn’t be made publicly available. Finally, the Docker daemon, controlled either through backgronud Celery jobs or directly by the `containers` Django app, spawns container instances for the user-created web apps. All containerized web apps run in a shared Docker network, isolated from the network where the Kiosc server and its supporting services run. Thus, the web apps can only by accessed through the reverse proxy implemented in the Kiosc web server.

### Operation

We provide a Docker Compose file which specifies all the necessary services and can be used to install and deploy Kiosc. Since Kiosc is intended for running interactive visualizations and not analysis workflows, it is designed to run on a single server and does not support horizontal scaling. This limits the number of containers that can run concurrently, but we found that the hardware resource usage also depends on the footprint of the individual web apps and how frequently they are visited. Because of the multiple apps and background jobs which will run concurrently, we recommend at least 4 CPUs and 8 GB of RAM, to be scaled up depending on the expected number of apps and users of the instance. Some apps are known to load large datasets in memory, thus hosting them on Kiosc will require larger machines. Our main instance, which was started in 2021, hosts 150 projects and 321 containers, of which 19 are running at the time of writing (Figure 2). The server is equipped with 16 Intel Xeon (Cascadelake) processors and 125 GB of RAM.

**Figure 2.**
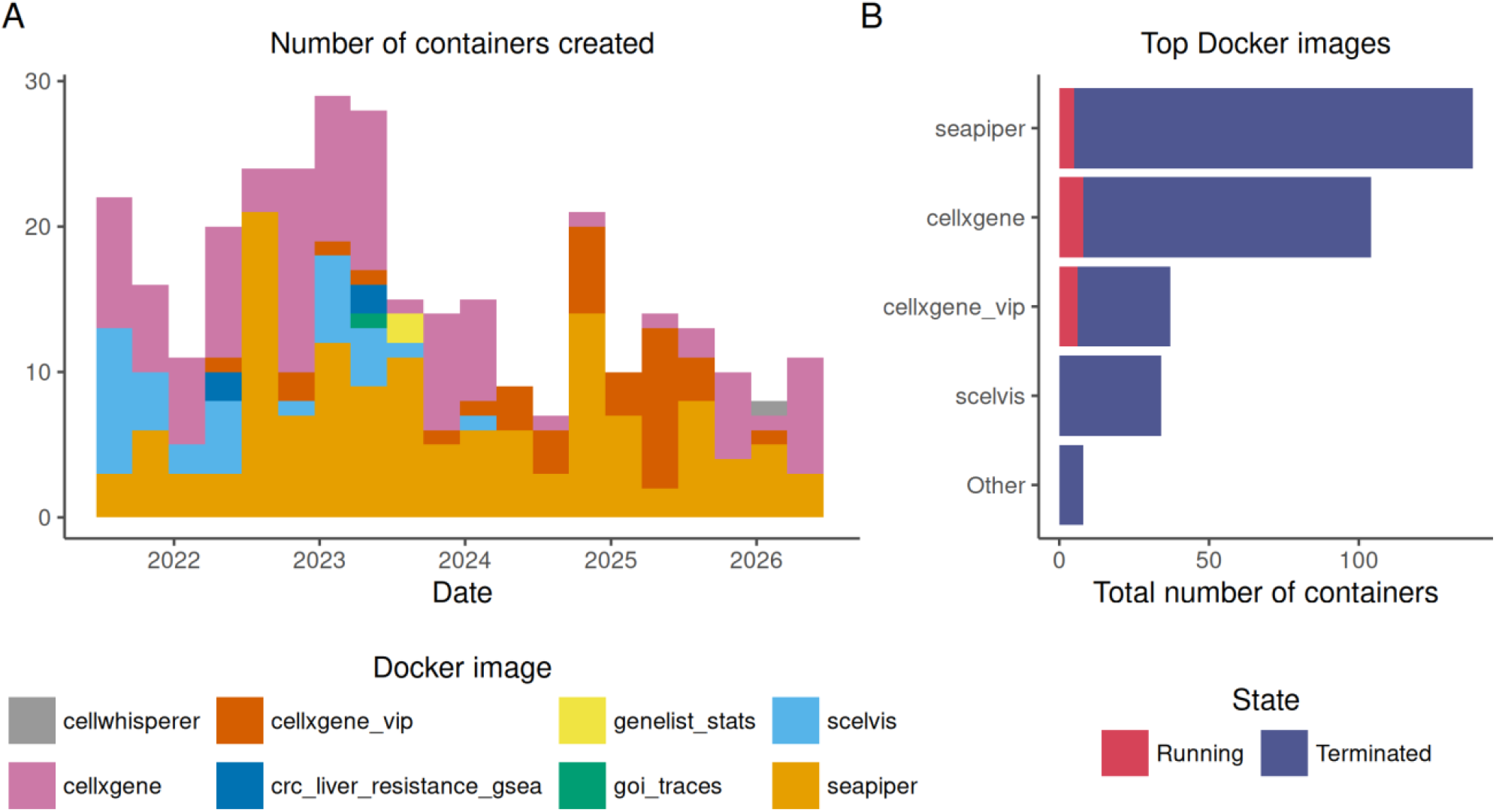
Summary of containers hosted in our internal Kiosc instance to date. (A) Histogram of containers created over time, colored by the underlying Docker image; each bin is 3 months wide. (B) Total count of container instances for the most popular Docker images, colored by their current state (either running or terminated).

There are three main types of users who interact with Kiosc. First, we have the instance administrator, whose role is to install the software at the site, keep it up to date, and troubleshoot when issues arise. Compared to alternatives such as Kubernetes, Kiosc is considerably simpler to operate (Table 1). The entire setup can be accomplished in a few command lines, and the software can be configured by environment variables. The second type of user is responsible for web app development. They will work hands-on on a data analysis project, then develop a bespoke Docker container for a web app where the analysis can be showcased or replicated and finally pull the container into Kiosc. Unlike ShinyProxy and OpenOnDemand, Kiosc allows users to add new web apps dynamically and autonomously from a web interface (Figure 3). Last, but not least, the biggest group of users is not necessarily trained in software development and data analysis, but can understand and make use of plots, tables, and biological insights resulting from the analysis. From their point of view, accessing a Kiosc web app will be no different than accessing a regular website.

**Table 1.** Feature comparison between Kiosc and other systems with overlapping use cases described in the main text.

|  | Kiosc | ShinyProxy | Open<br>OnDemand | Kubernetes | DockGE |
| --- | --- | --- | --- | --- | --- |
| Access control | ✓ | ✓ | ✓ | ✓ | ✗ |
| Dynamic container creation | ✓ | ✗ | ✗ | ✓ | ✓ |
| Scalability and resource control | ✗ | ✓ | ✓ | ✓ | ✗ |
| Workflow support | ✗ | ✗ | ✓ | ✓ | ✗ |
| Intended infrastructure | Single machine | Single machine or cluster | Mostly cluster | Mostly cluster | Single machine |
| System set-up | Docker compose and environment variables | Docker and YAML file | System package (or ansible) and YAML file | Management binaries and YAML files | Docker compose |

**Figure 3.**
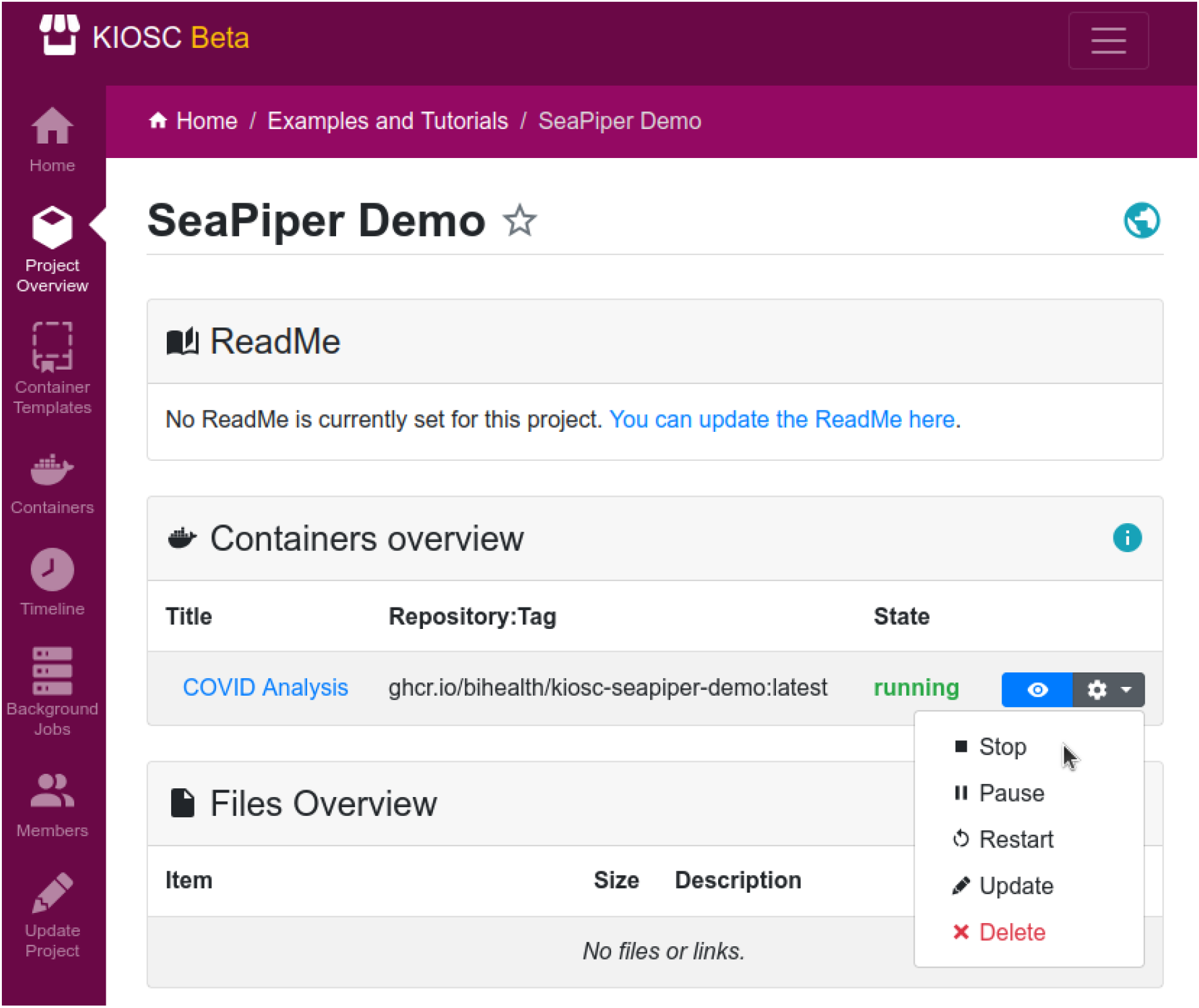
Screenshot from the Kiosc web interface. The Project Overview page features a list of the containers and files in the project, if any. Containers can be accessed and managed from this page.

A typical workflow is as follows. A user will first create a project and possibly add other users as members. Then, he or she will create a container object by filling a form with all the necessary details: a title for the container, the Docker image name and tag, the network port at which the web app can be reached, and optionally the environment varaibles that should be made available inside the container; it is also possible to override the default command that is launched when the container starts. The members of the project to which the container belongs can then start, stop, and access the web app. Optionally, a Kiosc instance can be linked to an upstream SODAR site (Nieminen et al. 2020), from which users and projects can be inherited.

Since Kiosc was designed with the goal in mind to share containers running a web app that presents data to users, containers often need to access data sets at runtime. While it would be technically possible to compile the data directly into a Docker container, it is not a recommended practice as it defies the purpose of a container to be reusable and would fill up the Kiosc Docker cache quickly. Kiosc offers multiple ways to use data within a container. First, smaller files can be uploaded directly from the web interface and stored in the PostgreSQL database. This method is not recommended for files in the gigabyte range or higher. Second, Kiosc can automatically download files from a URL (supporting multiple protocols, including HTTPS and FTPS) and save them into persistent Docker volumes, which are then mounted in the container at a user-defined location. Third, the creator of the Docker container can integrate the functionality to load data from an external source into the app once the container starts. This can be achieved either by setting environment variables that the web app can interpret (sensitive variables can be marked and the value will be then masked in the container setting), or, if the web app allows, by defining a custom command to be run when the container starts. Many web apps, including Shiny and CELLxGENE, are designed to read such parameters and load the data accordingly.

## Use Cases

### Fostering interdisciplinary collaboration in bioinformatics

In a field that increasingly relies on big data (Altaf-Ul-Amin et al., 2014; May, 2014; Pal et al., 2020; Sánchez-Tójar et al., 2025), we regard scientific web apps as a bridge between computational and wet-lab biologists, empowering the latter to make meaningful contributions in the data analysis process. Computational biologists develop the analysis using sophisticated statistical methods and produce easily interpretable figures, tables, and interactive widgets; wet-lab biologists and clinicians close the loop by providing interpretation and new questions to address, based on the analysis results. In our experience, this workflow is greatly simplified when all parties share the same platform where the results can be explored, reproduced, and iterated upon. Since Kiosc web apps can be made available through the internet and are subject to access regulation, it is easy and secure to share them among colleagues and selected collaborators, even across institutions.

In the past three years we initiated 46 projects which involved interactive exploration for single-cell transcriptomic data, using either SCelVis (Obermayer et al., 2020) or CELLxGENE (CZI Cell Science Program et al., 2025). As a bioinformatics core unit and research lab, we set up Docker containers with relevant data and uploaded them to our Kiosc instance. Our collaborators were then granted access to the apps and data.

Another recurrent task in our group consists of running bioinformatics pipelines for analyzing primary data. Since the output from these pipelines tends to be verbose, we developed our own apps to explore the results interactively. One of the most popular has been seaPiper (Weiner, 2025) (Figure 4), which we run in over 60 projects. SeaPiper is a Shiny app for visualizing the output of the SeA-SnaP pipeline, a Snakemake workflow for sequencing data analysis (https://github.com/bihealth/seasnap-pipeline).

**Figure 4.**
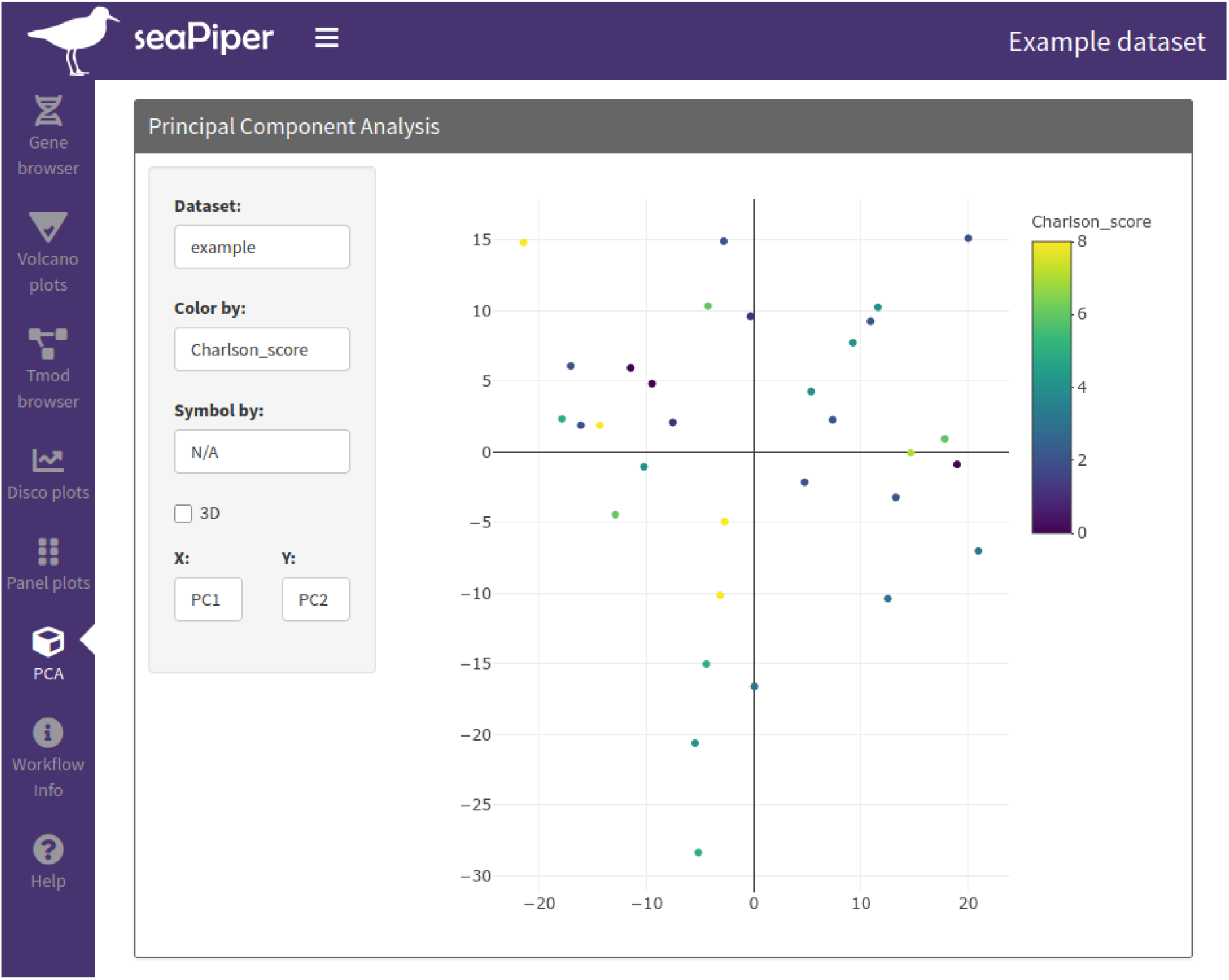
View of the seaPiper app proxied through Kiosc. From a user’s perspective, accessing an app from Kiosc looks and feels just like accessing a regular web site. The fact that the design of the Kiosc and seaPiper are similar is just a coincidence.

### Creating companion apps for scientific articles

Sharing the code used to obtain the results shown in an article increases the robustness and reproducibility of science (Laurinavichyute et al., 2022; Miske et al., 2026; Springer Nature, n.d.). Some authors even argue for the concept of an “executable paper”, where text, raw data, code, and results are combined in a single document that the reader can interact with (Lasser, 2020). Indeed, even if source code is available, it can be difficult to use on its own. On the other hand, if, along with the source code, the main figures and tables are also browsable with a simplified web interface, it becomes easier to find the relevant information. Kiosc can be used to publish apps accompanying articles, enabling users to reproduce the results from the paper or, if the app supports it, replicate the same analysis on their own data. In these cases, the container can be made publicly accessible through the internet. This use case would not be feasible just with Open OnDemand, which is rather meant to allow authorized users to access an institution’s compute resources. Kiosc, on the other hand, can be deployed in a safe and isolated machine.

### Making data exploration more reproducible

A common problem is that software which works today may stop functioning tomorrow. In Kiosc, web apps are packaged in Docker containers, which include fixed versions of the dependencies of the software, giving your app the best chances of being runnable several years into the future (Boettiger, 2015). Different projects, especially if started at different timepoints, may require different versions of the same app. Installing the apps on a personal or shared computing environment directly can lead to conflicts between dependencies. Kiosc can also solve this problem by isolating the apps inside containers and making them available simultaneously through its web interface. By organizing containers into projects and categories, Kiosc simplifies the management of many data analyses reports. When combined with computational notebook tools such as Jupyter (Granger & Perez, 2021) and Quarto (Allaire et al., 2022), Kiosc can be used to host fully reproducible and browsable project reports. If the notebooks perform heavy computations at runtime, the reports can be precompiled into static HTML files and just served through Kiosc, without having to rely on third-party solutions such as GitHub pages (https://pages.github.com).

### Teaching using engaging material

Finally, we believe Kiosc can also be used productively in the classroom. Here, the main advantage is the ability to create fixed and reproducible environments at scale, potentially using the Kiosc API for bulk operations. This makes sure that all students have access to the same experience without having to install the app on their own personal computers.

## Conclusions

Developing and using web apps in science provides a number of advantages, but their adoption can be hindered by lack of appropriate infrastructure to publish them and keep them available long after the end of the project or the publication of the corresponding article. Kiosc addresses this by providing an easy-to-use platform where web apps can be published with little maintenance overhead.

Packaging the apps in Docker containers also helps guarantee that the software will remain functional despite the instability of modern development ecosystems, which are subject to frequent version changes and fast turnover of technology stacks (Boettiger, 2015).

In this work, we focused on some of the most representative ways we have interacted with our own Kiosc instance over the years. However, the platform is not limited to the apps we described here. For example, in the past we have successfully configured ProTIGY (https://github.com/broadinstitute/protigy-v2), a toolset for proteomic data analysis. More broadly, the platform is compatible with any web app, so long as it is packaged in a Docker container.

For many of the use cases covered in this paper, it is often necessary to rapidly adopt new apps or update them to the latest version. This process can be greatly streamlined if users are allowed to do these tasks dynamically and autonomously, without the mediation of an administrator. This can be done with Kiosc but not with other hosting tools.

In the future, we plan to migrate to Kiosc some of the services that we maintain as a Bioinformatics Core Unit. For example, we support a local UCSC Genome Browser instance (Casper et al., 2026) to which custom tracks can be uploaded. Hosting this service in Kiosc instead of a permanently running dedicated virtual machine achieves higher flexibility, since users can start and stop their own instances depending on their needs, freeing resources when the usage demand is low.

Another planned development is to upgrade the hardware of our Kiosc instance and equip it with a graphics card. This will allow apps to render plots faster if they use specialized libraries like OpenGL. On the other hand, it will also enable Kiosc to support modern tools based on locally-hosted large language models such as CellWhisperer (Schaefer et al., 2025), an artificial intelligence (AI) model and software tool for chat-based interrogation of gene expression datasets. One of the biggest limitations of Kiosc is that currently containers can only run on a single machine, therefore they will compete for the same resources. Besides upgrading our hardware, we will also implement a solution to distribute containers across machines, potentially using Docker Swarm mode.

By developing Kiosc, a low-maintenance platform for publishing web apps, we hope to encourage collaboration and increase the transparency of bioinformatics research.

## Data availability

Kiosc is open source software licensed under the MIT License. The source code is available from: https://github.com/bihealth/kiosc-server. Data which is part of the scientific web apps hosted on Kiosc is available on the platform.

## Author contributions

- Federico Marotta: Software, Writing—Original Draft, Investigation, Visualization.
- Oliver Stolpe: Software, Writing —Original Draft, Investigation, Visualization.
- Benedikt Obermayer: Software, Investigation.
- January Weiner: Software, Investigation.
- Manuel Holtgrewe: Conceptualization, Software, Supervision, Writing —Review & Editing.
- Dieter Beule: Conceptualization, Funding acquisition, Resources, Project Administration, Supervision, Writing —Review & Editing.
- Mikko Nieminen: Conceptualization, Software, Visualization, Supervision,Writing— Review & Editing.

The manuscript was seen and approved by all authors and it hasn’t been accepted or published elsewhere.

## Acknowledgements

We are thankful to the whole AG Beule team, the Core Unit Bioinformatics (CUBI), and especially Eric Blanc for helpful feedback, beta-testing, and adoption of Kiosc.

---

**Figure S1.**
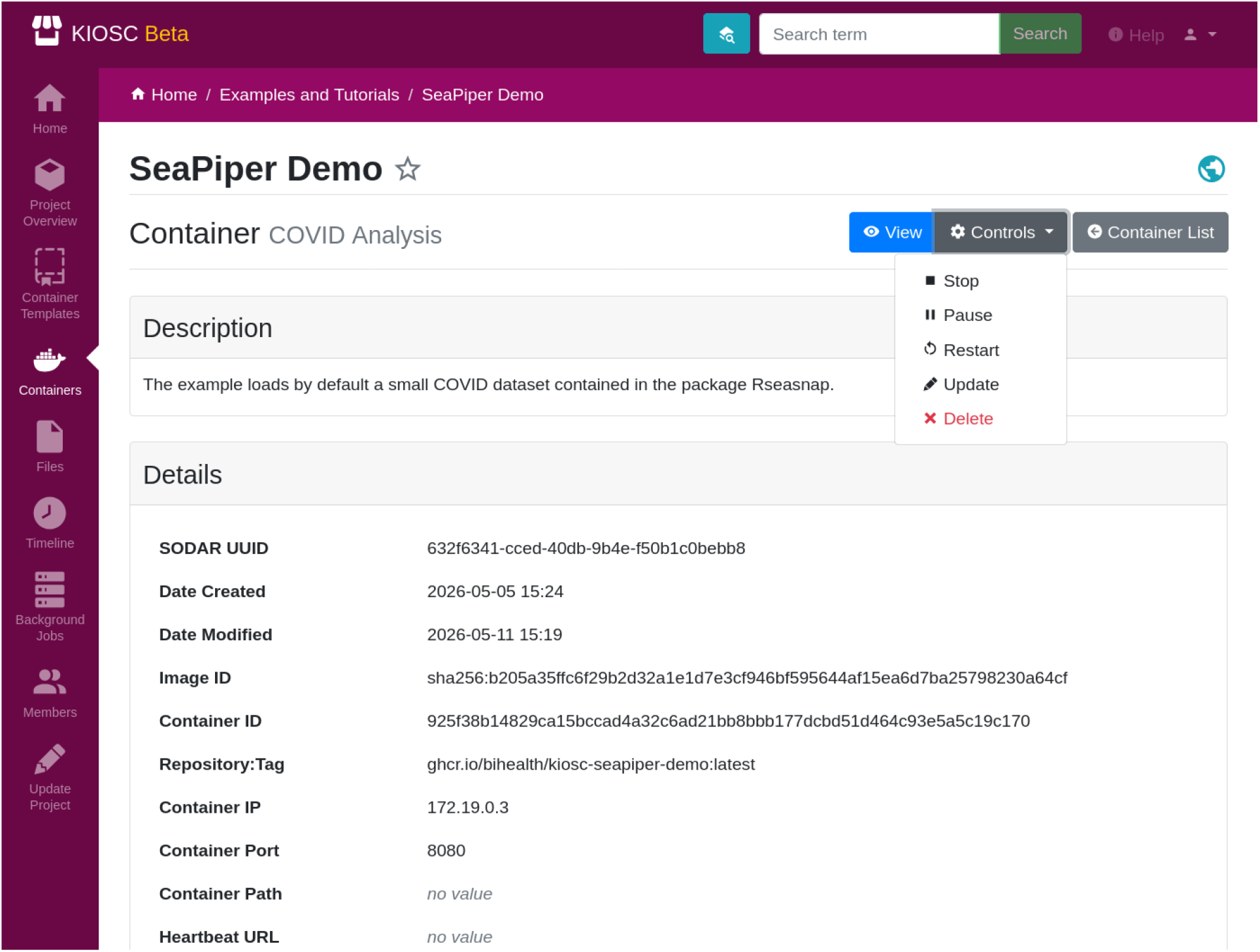
Additional screenshot from the Kiosc user interface. The “Container Detail” view summarizes the technical properties of the Docker container. The left side menu also shows six of the eight Django apps as mentioned in the Mehods section, among others: `containertemplates`, `containers` from Kiosc itself and `projectroles`, `filesfolders`, `timeline`, `bgjobs` from SODAR Core.

**Figure S2.**
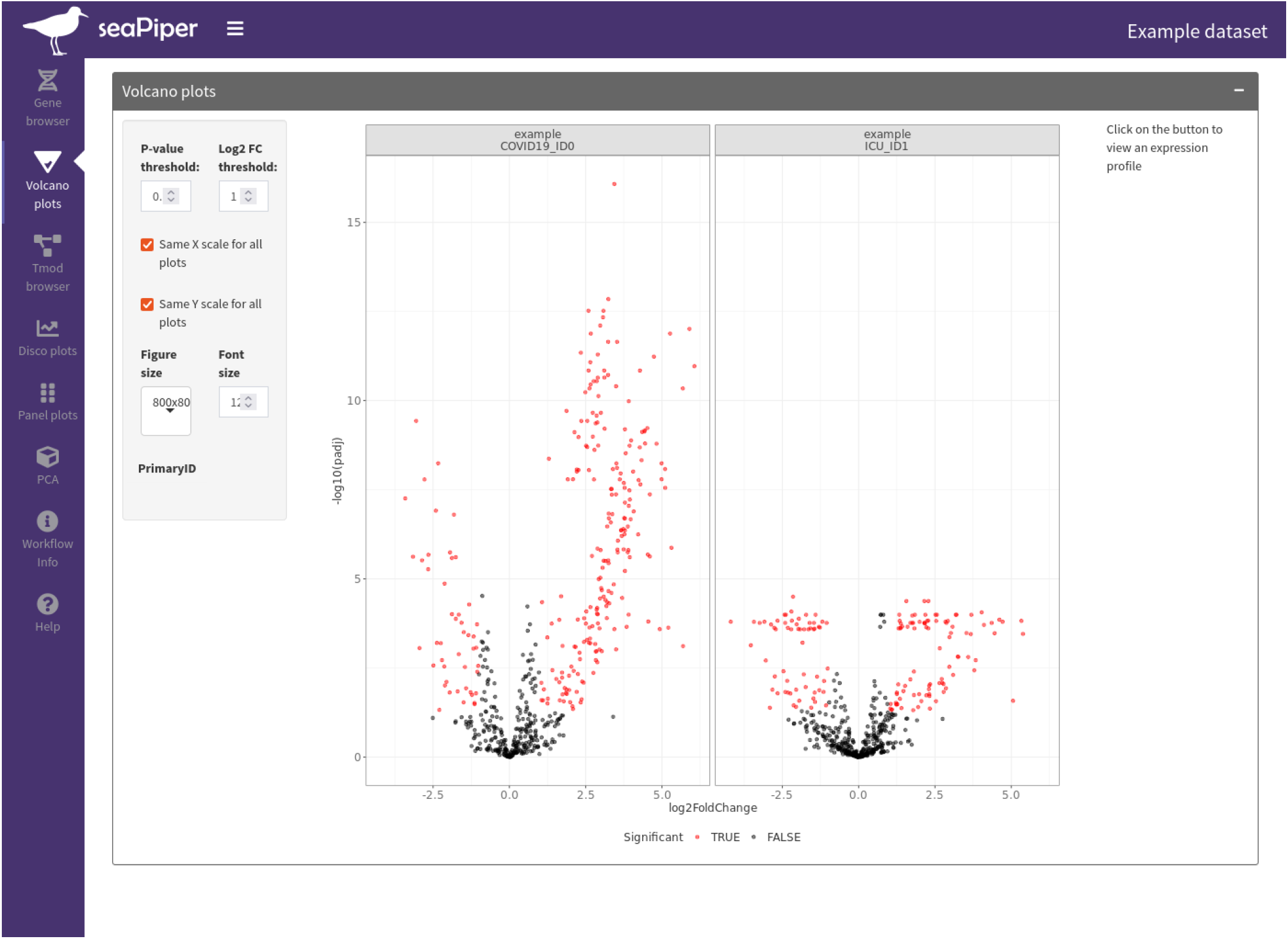
Additional screenshot from the seaPiper app.

